# Sparse memory ensembles set brain-wide network states to sustain learned associations

**DOI:** 10.1101/2025.03.07.642018

**Authors:** Josué Haubrich, Gabriele Russo, Denise Manahan-Vaughan

## Abstract

The ability to recall spatial memories and adapt behavior to changing conditions is essential for efficient navigation of environments. These processes are thought to rely on sparse populations of neurons, embedded within distributed brain networks. However, how the activity of these neurons impacts large-scale network connectivity to sustain behavior remains poorly understood. To address this, we tagged neurons active during peak performance in an appetitive spatial memory T-maze task in male and female TetTag-hM3Dq transgenic mice and chemogenetically reactivated these ensembles during functional magnetic imaging (fMRI) of brain activity. Reactivation of these ensembles triggered widespread network reorganization, optimizing the brain’s modular specialization while maintaining integration across the network via key hubs and gateways. We identified two distinct clusters dominated by the ventral and dorsal hippocampus, respectively, that exhibit distinct connectivity with cortical and subcortical areas. Control mice showed effective extinction learning (EL) in the T-maze when the reward incentive was absent. Reactivating the tagged memory ensembles during EL significantly prevented this process. These findings reveal the brain-wide network hubs that support effective spatial memory acquisition, but also indicate that decay of this network connectivity is likely to be an essential facet of effective EL of spatial experience.

## INTRODUCTION

Navigating environments, and adapting strategies to changing contingencies, rely on activity in specific brain structures, as well as on complex, brain-wide network interactions (Ekstrom et al., 2017; Henson and Gagnepain, 2010). Memories of these learned experiences are encoded by sparse populations of neurons distributed across interconnected circuits (Josselyn and Frankland, 2018). Although manipulating these sparse memory ensembles within local circuits can trigger or block the expression of consolidated memories (Josselyn et al., 2015), it remains unclear how the activity of these ensembles can drive brain-wide information flow and system dynamics (Goode et al., 2020). Here, we investigated how the reactivation of neurons, that were engaged during the retrieval of a well-learned appetitive spatial memory task, influence memory during spatial appetitive extinction learning (EL). We also examined how brain connectivity is represented, by using functional magnetic resonance imaging (fMRI), during this reactivation event,

The dorsal hippocampus is essential for spatial representation and episodic memory (Eichenbaum et al., 1999; Manahan-Vaughan, 2017), providing a spatial-temporal framework that enables the creation of a ’cognitive map’ of the experienced world (Stachenfeld et al., 2017). It is particularly critical for encoding trajectories toward rewards (Frank et al., 2000), integrating goal representations within a what-when-where framework (Montagrin et al., 2024), pattern completion and pattern separation (Lee et al., 2020; Maurer and Nadel, 2021), as well as supporting memory formation, recall, and updating (Coelho et al., 2024; Haubrich et al., 2017, 2015; Lacagnina et al., 2019; Sekeres et al., 2017). Also essential for effective navigation in a changing world, is the ability to recognize that a previous association no longer leads to an expected outcome during EL (Bouton et al., 2021; de Oliveira Alvares and Do-Monte, 2021), a process that also engages the dorsal hippocampus (Mendez et al., 2018; Méndez-Couz et al., 2019; Pignatelli et al., 2019), as well as the prefrontal cortex (PFC) (Quirk and Mueller, 2008).

Although it was previously thought to support non-spatial information processing, including anxiety-based behavior (Strange et al., 2014), more recently it has become apparent that the ventral hippocampus also supports spatial learning (Contreras et al., 2018; Lee et al., 2019; Ramos and Morón, 2022). The ventral hippocampus also plays a key role promoting goal-oriented learning and information updating (Avigan et al., 2020; Blanquat et al., 2013; Ruediger et al., 2012) that are supported through its interconnectivity with the PFC and dorsal hippocampus (Komorowski et al., 2013; Park et al., 2021).

Consolidated associative memories are stored in distributed brain structures beyond the hippocampus (Goode et al., 2020), with interregional interactions supporting encoding and retrieval (Teyler and Rudy, 2007; Tonegawa et al., 2015), behavioral modulation (Eichenbaum, 2017) and the updating of predictions during extinction learning (Bouton et al., 2021). The retrosplenial cortex (RSP) processes spatial, cognitive, and reinforcement information (Stacho and Manahan-Vaughan, 2022a), while the cingulate cortex (ACC) supports long-term memory storage (Frankland and Bontempi, 2005; Haubrich et al., 2016) and cognitive control (Shenhav et al., 2013). The posterior parietal cortex (PCC), by contrast, contributes to multimodal spatial processing and encoding (Kravitz et al., 2011). In the PFC, the orbitofrontal region (ORB) governs decision-making and association updating (Padoa-Schioppa and Conen, 2017; Ranganath and Ritchey, 2012), while the infralimbic (IL) region facilitates decision-making, response adaptation, and extinction learning (Quirk and Mueller, 2008; Roughley and Killcross, 2021). This contrasts with the actions of the prelimbic (PrL) region, that counteracts these processes (Howland et al., 2022; Nett and LaLumiere, 2021).

Integral components of brain networks, thalamic nuclei such as the paraventricular (PVT), reuniens (RE), and laterodorsal (LDT) relay sensory information (Roy et al., 2022), support memory functions (Arts et al., 2017; Chao et al., 2022), motivate behavior (Penzo and Gao, 2021), direct attention (Fiebelkorn and Kastner, 2020), and aid spatial navigation (Aggleton and O’Mara, 2022; Perry and Mitchell, 2019). Moreover, the nucleus accumbens (NAc) is critical for adaptive navigation since it associates environments with salient outcomes (Ito et al., 2008) and supports spatial appetitive learning and retrieval (Trouche et al., 2019).

While spatial navigation and memory involve numerous brain areas, these processes are thought to depend on the activity of sparse neuronal populations within each structure (Josselyn and Frankland, 2018; Sweis et al., 2021). In mice, combining long-lasting activity-dependent cell labeling with targeted manipulations has enabled the study of specific neuronal ensembles activated during aversive experience (Garner et al., 2012; Reijmers et al., 2007). This approach allowed gain- and loss-of-function studies to investigate memory-related ensembles across brain regions, including the hippocampus, amygdala, RSP, prefrontal cortex, and the NAc (Tonegawa et al., 2015). While reactivating or silencing tagged memory ensembles can promote or suppress learned behaviors, it remains unclear how the activity of these sparse populations influences brain-wide communication, the extent to which they might be engaged in non-aversive forms of learning, and whether the resulting systemic changes drive behavioral responses.

In this study, we hypothesized that memory-related neuronal ensembles across brain areas influence behavior not only by modulating local circuitry, but also by altering patterns of brain-wide information flow. We also tested the hypothesis that the reactivation of a well-acquired spatial memory, in the absence of a reward incentive, will disrupt EL of this task. We also assessed the contribution of neurons. Within brain-wide networks in this process.

Using TetTag-hM3Dq mice (Cai et al., 2016; Khalaf et al., 2018; Reijmers et al., 2007) to label and subsequently reactivate neurons that were engaged during successful completion of a spatial appetitive learning task, we assessed whether reactivating these neuronal ensembles could sustain learned behaviors without reward. By means of functional magnetic resonance imaging (fMRI) during reactivation, we also examined whether the activity of these neuronal populations alters interregional coordination and functional connectivity across the brain, reflecting the effective acquisition of a spatial appetitive task. Our findings reveal that reactivating these neurons during exposure to the same paradigm in the absence of an appetitive reward, prevents successful EL. Neuronal re-activation revealed of hub-like regions of brain connectivity that are likely to reflect the effective acquisition of spatial appetitive memory.

## METHODS

The study was carried out in accordance with the European Communities Council Directive of September 22nd, 2010 (2010/63/EU) for care of laboratory animals and all experiments were conducted according to the guidelines of the German Animal Protection Law. The study was approved, in advance, by the North Rhine-Westphalia (NRW) State Authority (Landesamt für Arbeitsschutz, Naturschutz, Umweltschutz und Verbraucherschutz, NRW). All efforts were made to minimize the number of animals used.

### Mice

We crossed heterozygous TetTag (cfos-tTA/cfos-shEGFP) and tetO-hM3Dq mice (both from The Jackson Laboratory, Bar Harbor, USA) to produce double transgenic TetTag-hM3Dq mice (TetTag group). Control animals were wild-type siblings (Wt group). Polymerase chain reaction (PCR) from ear punch tissue (obtained at <3 postnatal weeks) was used to detect the transgenes. The PCR primers for cfos-tTA/cfos-shEGFP were 5’ AAG TTC ATC TGC ACC ACC G 3’ and 5’TCC TTG AAG ATG GTG GTG CG 3’, and for and tetO-hM3Dq were 5’ CGT CAG ATC GCC TGG AGA 3’ and 5’ CGG TGG TAC CGT CTG GAG 3’ (biomers.net GmbH, Ulm, Germany).

We used 8-12 week-old male and female mice (25-35g), housed in individual cages, with ad libitum access to water, under a 12-hour light/dark cycle with controlled humidity and temperature. Animals were fed with food containing doxycycline (dox; 625 mg/kg; Ssniff, Soest, Germany) and were weighed before commencing the study and subsequently maintained at 85% of their initial body weight. Each mouse was handled daily for at least 5 days before the behavioral procedures. Both groups underwent identical experimental procedures. All experiments took place during the light phase.

### Treatments

Deschloroclozapine (DCZ; Tocris, Bristol, United Kingdom) was freshly prepared on the day of administration by dissolving it in 2% dimethyl sulfoxide (DMSO) and physiological saline under sonication. DCZ was administered via intraperitoneal injection (i.p.) at a dose of 0.1 mg/kg. Both Wt and TeTag animals received this treatment.

### T-Maze protocol

Experiments were conducted in a T-Maze consisting of a starting box made of gray polyvinyl chloride (22 x 10 cm), a central arm (90x 10 cm), a decision arm (100 x 10 cm), and two return arms (85 x 10 cm), with 19 cm high walls. Distal visual cues placed outside the maze included a cylinder with horizontal stripes to the left, a cylinder with vertical stripes to the right (both made of white cardboard, 50 cm high and 20 cm wide, located 12 cm distally from the respective central arm walls and 15 cm from the inner lateral walls). A white cardboard panel (50 x 50 cm) containing a black circle (25 cm diameter) was positioned 30 cm behind the central arm’s end. A vanilla scent was applied to the end of both decision arms that could only be detected when the animals were at the end of the arms (Wiescholleck et al., 2014). The maze was housed in a tent illuminated by white LEDs at 30 lux, and trials were recorded using EthoVision XT v14 (Noldus, Wageningen, The Netherlands).

Mice were pre-handled and subjected to mild food deprivation before training, and their weights were monitored to maintain 85% of their pre-diet weight. Habituation began with 3 daily sessions of 10 minutes in the starting box. This was followed by 6 habituation trials over two days, where mice explored the entire maze. During these trials, they were guided to make a right or left turn without backtracking through the main maze arm, i.e. returning to the starting box via the return arms. Food pellets (Dustless Precision Pellets #F05684, Bio-Serv, San Diego, USA) were scattered throughout the maze to encourage exploration.

The acquisition phase began the following week. Mice were placed in the starting box and performed 20 trials per day, to locate a pellet reward that was consistently positioned at the end of the same decision arm. If a preference for one arm was observed during habituation, the reward was placed in the opposite arm. Trials began with the opening of the door to the central arm. Upon reaching the end of a decision arm (left or right), mice were blocked from backtracking and redirected through the return arm to the starting box, marking the end of the trial. During the first three days of acquisition, one pellet was consistently available in the target (rewarded) arm. On acquisition days 4 and 5, the reward probability was progressively reduced from 100% to 40% every five trials, while the reward magnitude was increased from one pellet to four pellets. Two days after the final acquisition session, mice underwent an extinction learning (EL) session consisting of 20 trials with no rewards. The following day, a final EL session was conducted.

A correct choice was defined as reaching the end of the rewarded arm, during the acquisition phase, whereas turning into the opposite (unrewarded) arm was considered an incorrect choice. If a mouse failed to leave the starting box within 40 seconds after the door to the central arm was opened, this was also counted as an incorrect choice. Similarly, failing to reach the end of a decision arm within 2 minutes was recorded as an incorrect choice. The percentage of correct choices was calculated for each block of 10 trials.

### Experimental design

TetTag mice were maintained on a dox diet to keep the TetTag system inactive. Wt animals received the same diet. The animals underwent training in a spatial appetitive task in the abovementioned T-maze (Figure 1) to learn to locate a reward that was placed with decreasing probability at the end of one specific T-maze arm (Acquisition phase, see above). After four days of training (days 1-4), consisting of two blocks of 10 trials each, and confirmation of successful task acquisition, animals’ rested undisturbed in their homecages. During this time, dietary dox levels were progressively reduced. By the end of the fourth day, the dox dosage was halved, and by the fifth day, dox was completely absent from the animals’ diet, opening, in the following day, a tagging window that allowed highly active neurons - i.e., those with high cFos expression - to express excitatory hM3Dq receptors (Reijmers et al., 2007). Because wild-type littermates lack the TetTag-hM3Dq transgenes, no tagging occurred in the control group, although controls also followed the same dox diet and dox withdrawal procedures.

**Figure 1.**
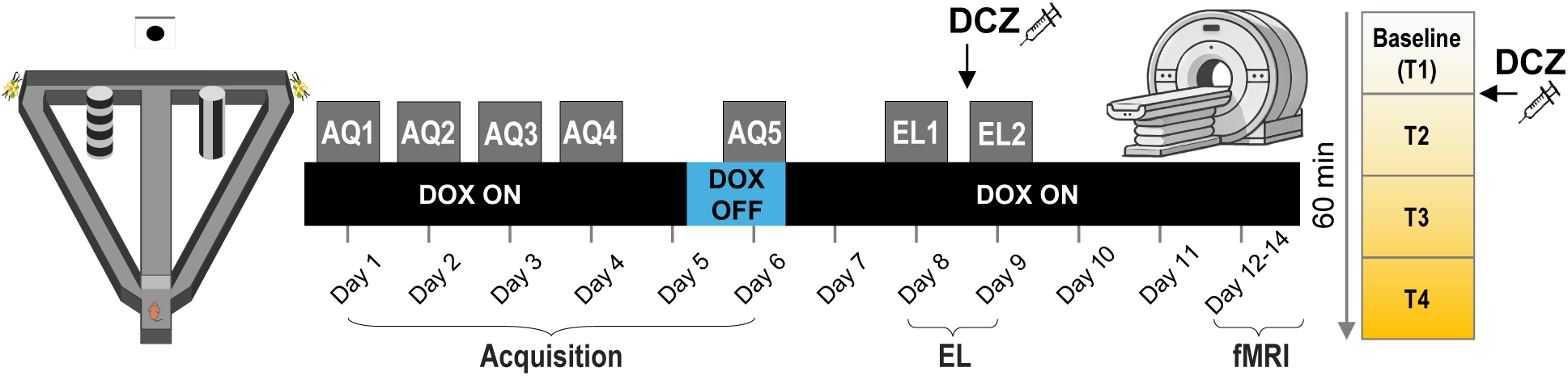
**Experimental design** TetTag-hM3Dq and Wt mice were trained in a T-maze task (left) to locate a low probability reward based on spatial cues (Acquisition phase: AQ1-AQ5). After four days of training, DOX was removed to open a tagging window, allowing neurons active during the final training session to express hM3Dq receptors. DOX was reintroduced to the animals’ diet, and two days later, mice underwent an extinction learning session without food reward (EL1). On day 9, a 2^nd^ EL session took place (EL2), where DCZ treatment occurred 30 min beforehand. Functional MRI scanning was performed three to five days later, in the presence of DCZ to assess brain-wide effects of neuronal reactivation.

On day 6, a final acquisition training session was conducted to tag neurons that were active during spatial memory retrieval and dox was reintroduced into the diet immediately afterwards (Figure 1). After micerested for two days in their homecages, EL was tested. This was conducted in the same context (AA paradigm) in the absence of any food reward, a protocol that requires maze exposure on two consecutive days for EL to be effective (Méndez-Couz et al., 2021). Thus, on day 8, EL sessions were begun, but on day 9, the animals were systemically injected with DCZ, to reactivate the cells labeled during the acquisition phase of the memory task (Nagai et al., 2020). DCZ is a potent, metabolically stable DREADD agonist, with negligible affinity for endogenous receptors, therefore its action is selective to neurons tagged with DREADD receptors (Nagai et al., 2020). The EL session on day 9 was commenced 30 min after DCZ administration. Between three and five days later, the mice were anesthetized for assessment using fMRI. After 15 minutes of baseline image acquisition, DCZ was administered, and scanning continued for an additional 45 minutes. (Figure 1).

### Immunohistochemistry

Immunohistochemistry was conducted post-mortem to verify activity-induced labeling by the TetTag-DREADD system. Animals were maintained on a dox diet. Prior to final acquisition training, one group of TetTag mice had dox removed from their diet, as described above. The other group continued to receive daily dox. Animals then engaged in a further acquisition session in the T-Maze so that hM3Dq receptors were inserted into neurons that expressed cFos during learning in the dox-free group. Two days later animals were injected with DCZ and 90 min later, they were perfused transcardially with 4% paraformaldehyde (PFA). Frozen 30 µm sections were prepared using as freezing microtome Cuttec S (Slee medical GmbH, Nieder-Olm, Germany). Sections were first washed thrice in PBS for 10 minutes each. A blocking solution containing 10% normal goat serum (n-Goat; Biozol Histoprime, Germany) and 0.2% PBS-Triton X-100 (PBS-Tx) was then applied, and the sections were incubated, during gentle shaking (Polymax 1040, Heidolph GmbH, Schwabach, Germany), for 90 min at room temperature. A primary antibody solution was prepared, containing rabbit anti-GFP (1:1000; Invitrogen) and guinea pig anti-c-fos (1:2000, Synaptic Systems, Goettingen, Germany) in 1% n-Goat and 0.2% PBS-Tx., Sections were incubated overnight at room temperature during gentle shaking. On the second day, sections were washed three times in PBS for 10 min each. A secondary antibody solution was prepared, containing goat anti-rabbit Cy3 (1:250; Biozol Diagnostica Vertrieb GmbH Hamburg, Germany) and donkey anti-guinea pig Cy5 (1:500; Biozol Diagnostica Vertrieb GmbH, Hamburg, Germany) in 1% n-Goat and 0.2% PBS-Tx, and sections were incubated for 90 min at room temperature with gentle shaking. After incubation, sections were washed thrice with PBS for 10 min each, mounted onto gelatin-coated slides, and left to dry overnight. The following day, slides were briefly rinsed with distilled water and covered with Dianova Mounting Medium with 4’,6-diamidino-2-phenylindole (DAPI) (immunoSelect®; Biozol Diagnostica Vertrieb GmbH, Hamburg, Germany). DAPI-Labeled sections were scanned (Axio SlideScanner) and images were analyzed using imaging software (ZEN, Carl Zeiss, Oberkochen, Germany) or QuPath (v 0.4.3)(Bankhead et al., 2017) to identify neuronal labeling.

### MRI and fMRI procedures

Imaging was performed using a small animal MRI system (7-Tesla horizontal bore, BioSpec 70/30 USR, Bruker, Germany), equipped with a cryogenically cooled quadrature surface coil for mouse heads (MRI CryoProbe, Bruker, Germany). Mice were anesthetized with isoflurane (3% for induction and 1.5% for maintenance) and stabilized on an animal bed by means of a conventional three-point fixation method that used a tooth-bar and earplugs (Z113760 and Z113768, Bruker, Germany) (Russo et al., 2021). Throughout the imaging sessions, body temperature was assessed via a rectal thermometer. A pressure sensitive pad placed underneath the abdomen monitored respiration (Small Animal Instruments Inc., New York, NY, United States). Monitoring of blood oxygen saturation and heart rate was conducted with a pulse oximeter clipped to the hind paw (Small Animal Instruments Inc., New York, NY, United States). Cooling and heating were enabled via water pipelines integrated into the mouse bed. Body temperature was maintained at 35.5 ± 1.5 °C during scanning. Imaging data was only collected at blood oxygen saturation values of 97–99%. At the beginning of each imaging session, a fast low angle shot (FLASH) localizer was used to roughly assess the position of the animal’s brain, then images in three different and manually adjusted planes were acquired one after the other using a multi slice FLASH sequence with the following parameters: repetition time (TR) = 522 ms, echo time (TE) = 3.51 ms, no average, acquisition matrix = 120 x 100, field of view (FOV) = 12 x 20 mm, spatial resolution = 0.1 x 0.2 mm^2^, slice thickness = 0.5 mm, number of slices = 16; subsequently, after the measurement of the map shim, a localized shim of the area of interest (whole brain) was performed.

On the basis of the sagittal reference images, 20 coronal slices (Supplementary Figure 1) were positioned along the entire brain and acquired with single-shot, multi-slice, gradient-recalled echo Planar Imaging (GRE-EPI), with the following parameters: Echo time (TE): 15 ms, TR: 1000 ms, slice thickness: 0.55 mm, no slice gap, matrix size: 95 x 60, FOV: 19 x 12 mm^2^, spatial resolution: 0.2 x 0.2 x 0.55 mm^3^, Bandwidth: 263 kHz, 4500 volumes. The total scan time was 60 min. The first 15 minutes (T1) served as a baseline period. At the end of T1, all animals received DCZ while remaining inside the MRI system. DCZ was administered intraperitoneally using plastic needles connected to saline-filled tubing, with an air bubble separating the physiological saline and DCZ. Following DCZ administration, scanning continued for an additional 45 minutes, divided into three bins of 15 minutes each (T2-T4).

At the end of the experimental session, high-resolution anatomical images, exactly aligned with the functional ones, were acquired using a RARE (**r**apid **a**cquisition with **r**elaxation **e**nhancement) sequence with the following parameters: TR = 2500 ms, effective TE (TEeff) = 15.55 ms, RARE factor = 4, number of averages = 4, FOV = 19 x 12 mm^2^, 20 coronal slices, slice thickness 0.55 mm, no slice gap, matrix size = 190 x 120, spatial resolution = 0.1 x0.1 mm^2^.

### fMRI data pre-processing and time-series extraction

Functional MRI data were processed in their native resolution using a standard pipeline (Grandjean et al., 2020) (Supplementary Figure 2). First, temporal spikes were removed using analysis of functional images (AFNI) (https://afni.nimh.nih.gov) 3dDespike. Subsequently, motion correction was performed (3dvolreg). Brain masks (BET, FMRIB Software Library, FSL)(Smith, 2002) were estimated on temporally averaged echo-planar imaging (EPI) volume (fslmaths). Then, spatial smoothing was applied using an isotropic 0.3 mm kernel (3dBlurInMask) corresponding to 1.5x voxel spacing along the dimension of the lowest resolution. Finally, bandpass filtering (0.01–0.1Hz) was applied (3dBandpass).

For normalization, the denoised and filtered individual scans were registered to reference space (Hikishima et al., 2017) using corresponding individual anatomical scans to enhance registration accuracy. Regions of interest (ROIs) were defined bilaterally in the reference space as 0.3mm^3^ spheres. The mean blood oxygen level-dependent (BOLD) signal time-series within a ROI were extracted (fslmeants) and subsequently utilized in the network analysis.**Network analysis**

### Network generation

We computed Spearman correlations between BOLD signals of all pairs of brain regions acquired during fMRI. Networks were generated by thresholding the interregional correlations for each animal and time bin. Only correlations exhibiting a Spearman p-value of below 0.001, after Bonferroni adjustment for multiple comparisons, were retained. Brain structures were defined as nodes in the network, and the corresponding r-values of the significant correlations served as the weights of the edges. Only positive correlations were considered. Networks were generated for each individual scan and time bin.

For analyses involving averaged networks, we retained correlations that were significant in at least 75% of individual networks within each group. These retained correlations were then averaged and considered as edges in the network. Community detection within these networks was performed using the Louvain algorithm (Blondel et al., 2008).

### Centrality measures

Nodal and global network measures were computed using the igraph package in R and custom R code (github.com/johaubrich) to evaluate properties of the functional networks. The metrics included:

- Global efficiency, the average of the inverse shortest path lengths between all pairs of nodes;
- Clustering coefficient, the ratio of the number of triangles to the number of connected triples in the graph;
- Edge density, the ratio of the number of edges present in the graph to the maximum possible number of edges between all pairs of nodes;
- Eigenvector centrality (Hub score), the values of the first eigenvector of the graph adjacency matrix, evaluating a node’s importance by accounting for its connections to other highly influential nodes;
- Betweenness, the number of shortest paths going through a node.

For centrality comparisons that were not based on averaged matrices, measures were averaged per group and time bin and expressed as a percentage of the mean value calculated during the baseline (T1). Differences in centrality measures between groups were compared by resampling the data with replacement 1,000 times, to generate bootstrap distributions of the mean differences between groups for each measure and time bin. We calculated 95%, 99%, and 99.9% confidence intervals using the bias-corrected and accelerated (BCa) method. Significance was determined based on whether these confidence intervals excluded zero.

We calculated the diversity score based on Shannon entropy (Shannon, 1948) to assess how many distinct communities were reached by each node’s neighbors. For each node, the communities of its neighboring nodes were identified, and the probability distribution of these communities was computed. The Shannon entropy of this distribution quantified “community diversity,” with higher values indicating that a node connects to a broader array of communities. Diversity scores were compared using the Wilcoxon test.

The Hub score was calculated by subtracting the nodal eigenvector centrality values of the Wt group from those of the TetTag group. To assess the significance of these differences, we performed a permutation-based test using network rewiring, generating null models through 1,000 iterations. P-values were computed by determining the proportion of times the absolute value of the differences from the rewired networks was greater than or equal to the absolute value of the observed difference.

To evaluate the effect of removing a node on the network’s global efficiency (GE), we computed the global efficiency of the network before node deletion minus its efficiency after deletion. Considering that brain inactivation has functional effects spreading to other connected regions (Grayson et al., 2016; Otchy et al., 2015), we employed a previously described disruption propagation model (Vetere et al., 2017).

### Statistics

For normally distributed data passing the Shapiro-Wilk test of normality, differences between groups were tested using the Student’s t-test. For non-normally distributed data, we employed the Wilcoxon test, bootstrapping, or permutation non-parametric approaches, as described above. Except for the generation of interregional correlation matrices, all other correlations were conducted using Pearson’s correlation test. All statistical analyses and plots were generated using R version 4.2.2 (R Foundation for Statistical Computing, Vienna, Austria).

## RESULTS

### Reactivation of spatial memory ensembles sustains previously acquired behavior during extinction learning

We first investigated how the reactivation of a successfully learned spatial appetitive learning task affects EL. TetTag-hM3Dq (TetTag) mice, and their Wt littermates, were maintained on a constant doxycycline (dox) diet. During a 5-day acquisition phase, the mice learned to locate a low-probability reward in a fixed location in a T-maze, relative to spatial cues. One day before the final acquisition session on day 6, dox was removed from the diet to open a window to tag neurons of TetTag mice, that were active during this session, by means of hM3Dq insertion. The animals then returned to the dox diet, and two days later, on days 8 and 9, underwent a 2-day EL phase. Thirty minutes before the second EL day, the animals received i.p. injections of DCZ to reactivate the tagged neurons in the TetTag group.

During the acquisition phase, no differences in performance were observed between TetTag and Wt mice across the training days (Mann-Whitney test, p > 0.05). Both groups showed significant improvement in correct choices from day 1 to day 3 and onwards (Wilcoxon signed-rank test, p < 0.05). Performance peaked on day 4 and remained stable on day 6 (the day of neuronal tagging) (Wilcoxon signed-rank test, p > 0.05) (Figure 2A, left).

**Figure 2.**
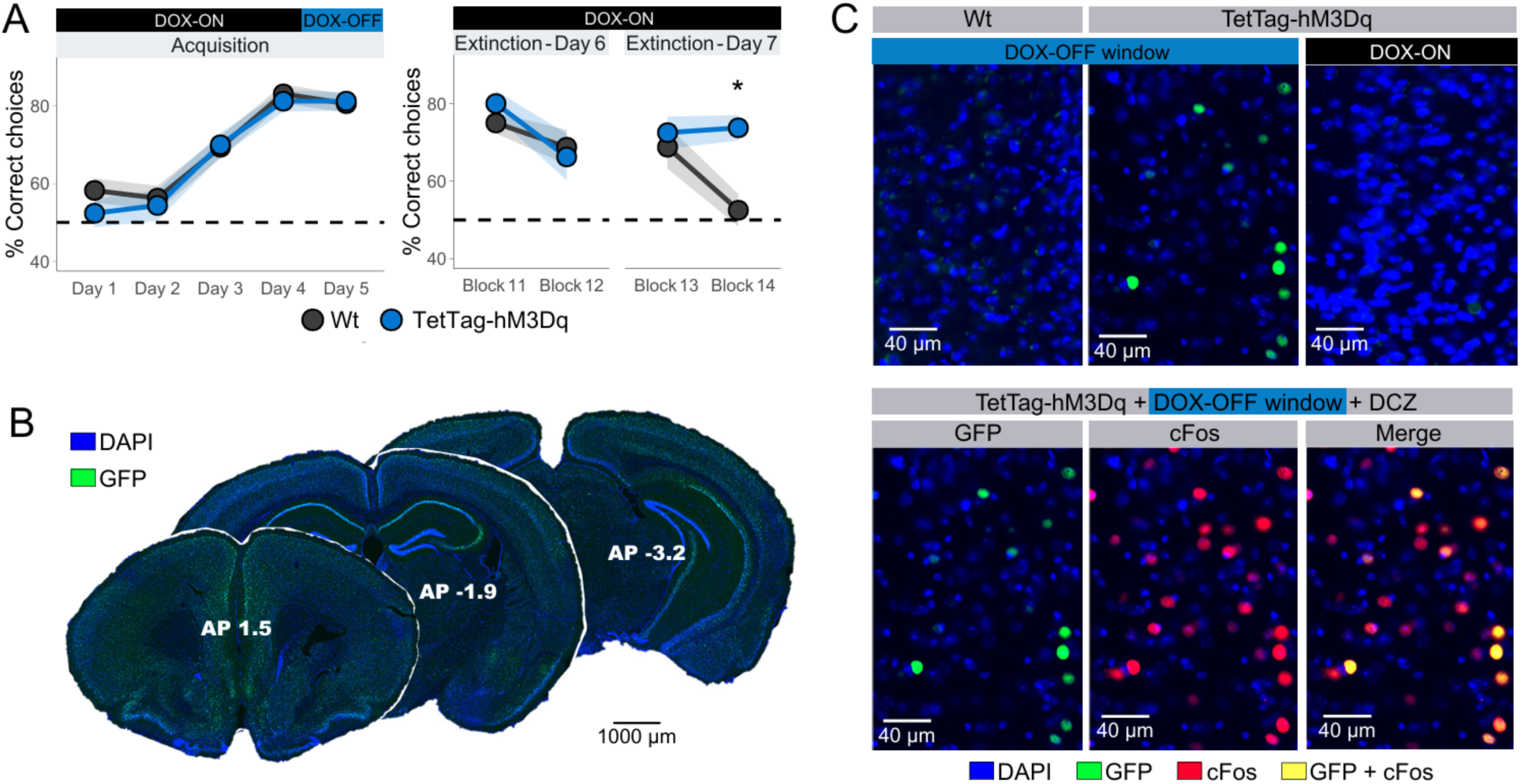
**Reactivation of Spatial Memory Ensembles Prevents Extinction Learning** A) Data points represent group means, and the shaded ribbon indicates the standard error of the mean. The upper bars denote the presence (black) or absence (blue) of doxycycline in the animals’ diet. N = 8 per group, *p < 0.05, Wilcoxon test. B) Representative low-magnification images of a TetTag animal after labeling during a dox-off window, showing sparse and widespread GFP-positive neurons. C)Representative coronal slices from a Wt and TetTag-hM3Dq mice showing tagged cells expressing GFP (green) co-stained with DAPI (blue) following either a dox-off window or a constant dox-on condition (top), and the effect of DCZ on the stimulation of labeled cells, verified by co-expression with cfos (red; bottom).

During the EL sessions, Wt mice exhibited a significant decrease in correct responses from the first through the final EL trial block (Wilcoxon signed-rank test, p < 0.05). Hence, treatment of the animals with DCZ 30 min before the second EL session did not affect EL in Wt mice. Both TetTag and Wt groups exhibited an identical correct choice behavior in the first trial block on day 9 (p > 0.05), that was not significantly different from their performance during the last trial block on day 8 (p > 0.05). However, the correct choice levels in DCZ-treated TetTag mice remained unchanged in trial block 14, indicating a failure to engage in EL as a result of the activation of the neurons involved in task acquisition (p > 0.05). A significant group difference was also observed during the final EL trial block on day 9 (Mann-Whitney test, p = 0.003; estimate = −20, CI (confidence intervals) [−30, −10]) (Figure 2A, right).

These results demonstrate that both Wt and TetTag mice developed a preference for the baited arm during the acquisition phase, achieving peak performance by the time the “dox-off” tagging window was open. During extinction learning, Wt mice required four blocks of 10 trials to extinguish the learned response. In contrast, reactivation of tagged neuronal ensembles via DCZ treatment in TetTag mice completely prevented extinction learning. These findings show that the reactivation of neuronal ensembles associated with spatial memory recall is sufficient to sustain learned responses, despite the absence of reinforcement during extinction learning.

Immunohistochemistry was used to verify effective neuronal tagging in TetTag mice, as well as DREADD-mediated neuronal stimulation (see methods). For this, GFP-expressing neurons were assessed. The dox-off period induced GFP-labeling in TetTag animals but not in wild-type (Wt) animals. TetTag animals maintained on a constant dox- on diet displayed no labeling (Figure 2B, 2C top). DCZ treatment reactivated the labeled neurons, as indicated by the co-expression of GFP and cFos (Figure 2C, bottom). In accordance with previous reports (Liu et al., 2012; Reijmers et al., 2007), the dox-off window during acquisition in TetTag animals induced labeling of sparse neuronal populations throughout the brain (Figure 2B).

### Similar BOLD signal acquisition as in controls, but different patterns of interregional correlations emerge following reactivation of spatial memory ensembles

To identify brain areas and networks that are engaged during successful spatial memory acquisition, we treated the TetTag and Wt littermates with DCZ during fMRI. BOLD signals were acquired from 21 regions: the dorsal and ventral dentate gyrus (d/vDG), d/vCA3, and d/vCA1 regions of the hippocampus; the dorsal subiculum (SUBd) and the postsubiculum (PoS); the reuniens (RE), laterodorsal (LDT), and paraventricular (PVT) thalamic nuclei; the nucleus accumbens (NAc); the orbitofrontal cortex (ORB); the infralimbic (IL) and prelimbic (PrL) regions of the prefrontal cortex; the anterior, intermediate, and posterior retrosplenial cortex (a/i/p RSP); the anterior and posterior anterior cingulate cortex (a/p ACC); and the posterior parietal cortex (PPC) (Figure 3A).

**Figure 3.**
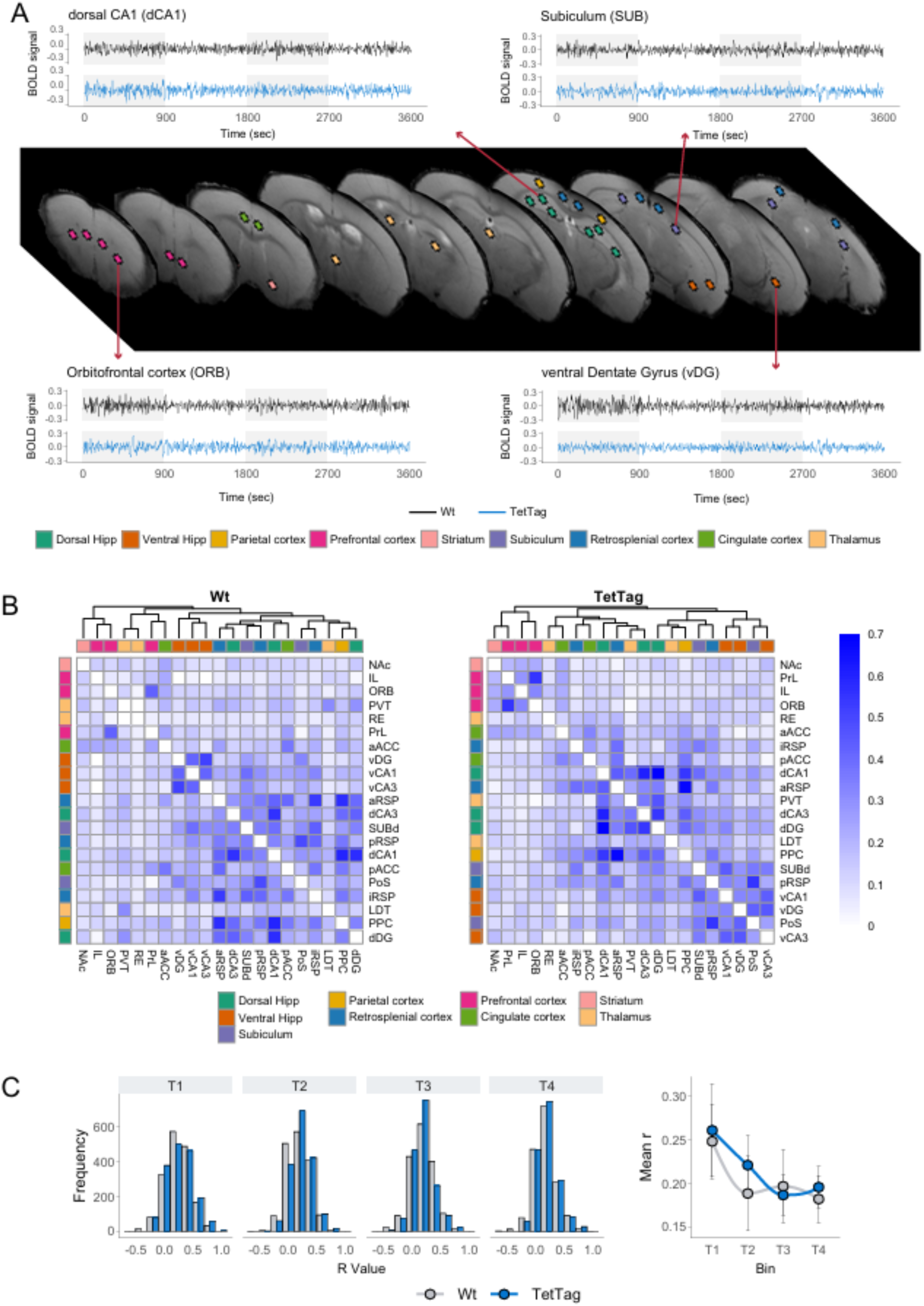
**Interregional correlations derived from BOLD signals.** A) Locations from where BOLD signals extracted and examples of signal time series from a Wt and a TetTag mouse over 60 minutes of scanning. B) Correlation matrices showing interregional BOLD signal correlations from the T4 bin for the Wt group (left) and TetTag group (right). Rows and columns are ordered by hierarchical clustering using the complete linkage method. Cells are color-coded according to Spearman’s correlation coefficient values. C) Top: Distribution of Spearman’s correlation coefficients (r values) across time bins. Bottom: Mean r values plotted across bins. N = 8 animals per group.

**Figure 4.**
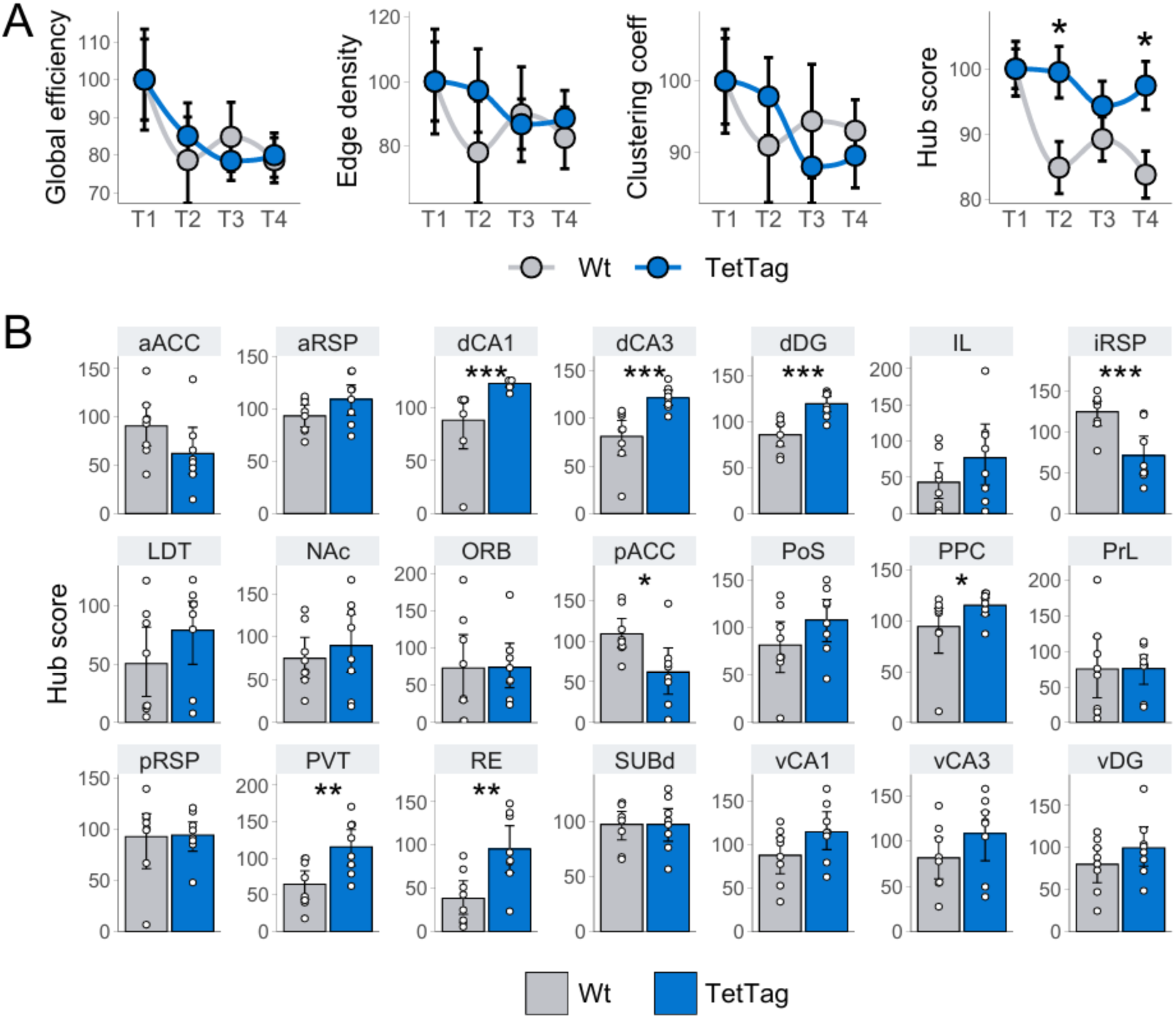
**Analysis of network centrality reveals increased Hub regions following reactivation** A) Network-level metrics of global efficiency, edge density, and eigencentrality (Hub score), across time bins and normalized to baseline values (T1). These metrics illustrate changes in network organization following memory ensemble reactivation at the end of T1. Points denote means and error bars the standard error of the mean. B) Hub score values of individual brain structures in the Wt and TetTag groups, indicating regions with altered network influence after engram reactivation in TetTag-hM3Dq mice. Bars denote means and error bars 95% confidence intervals. Asterisks denote significant differences between groups determined by bootstrapping. *p < 0.05 (95% CI), **p < 0.01 (99% CI), ***p < 0.001 (99.9% CI). N = 8 animals per group.

To assess the coordinated activity between brain structures, interregional Spearman correlations were calculated for each group and bin (Figure 3B). The analysis focused on T4, the timepoint corresponding to peak DCZ levels in the brain (Nagai et al., 2020; Nentwig et al., 2022) and the period corresponding to when behavioral test was conducted (Figure 1B). Reactivation of tagged memory ensembles in TetTag mice revealed interregional clustering patterns that were not evident in control mice (Figure 3B). However, the mean Spearman correlation coefficients across bins were similar between groups (Figure 3C), and no significant between-group differences in correlation values were found at any bin (t-test, p > 0.05), nor were there significant within-group changes across bins (paired-sample t-test, p > 0.05) (Figure 3C), showing that there were no changes in overall coordinate activity.

These findings indicate that reactivating the spatial memory ensembles that were engaged during memory acquisition does not alter overall levels of interregional correlations when responses in TetTag and control mice are compared. Rather the reactivation reveals the coordinated activity between specific brain structures that were presumably engaged during memory acquisition.

### Reactivation of spatial memory ensembles reveals functional network hubs

To further explore the interregional connectivity that emerged when the acquisition ensemble was reactivated, graph networks were constructed based on the computed correlations (Sporns, 2018). Brain regions were represented as nodes, while significant Spearman correlations between node pairs formed the edges. The weights of these edges corresponded to the correlation coefficients.

To assess the broad effects of spatial memory ensemble reactivation on brain connectivity, network centrality measures were calculated for individual graphs corresponding to each animal and time-bin, and normalized to each group’s baseline level (T1). We first calculated the network’s global efficiency (the ease of information transfer between nodes) (Vragovia et al., 2005), edge density (the ratio of observed connections to the total possible connections) (Wijk et al., 2010), and clustering coefficients (the density of triplets within the network) (Sporns et al., 2007). No significant differences between groups were observed across time points for these metrics (Figure 3A). We then calculated hub scores via eigenvector centrality (also referred as eigencentrality and prestige score), a metric that reflects the importance of a node by considering both its connectivity and the connectivity of its neighbors (Binnewijzend et al., 2013; Sporns et al., 2007). This is used to obtain insights into the emergence of highly influential nodes. The TetTag group exhibited significantly higher average hub scores compared to the Wt group at T2 (95% CI [3.52, 26.1]) and T4 (95% CI [2.33, 23.1]), indicating the formation of more prominent network hubs (Figure 3A).

To investigate whether specific structures gained prominence following spatial memory ensemble reactivation, hub scores of individual nodes were compared between groups (Figure 3B). Significant increases in hub scores were observed for dorsal hippocampal regions (dCA1, 99.9% CI [14.5, 85.1]; dCA3, 99.9% CI [17.2, 81.3]; and dDG, 99.9% CI [3.21, 58.8]), as well as for associative cortical and thalamic areas (posterior parietal cortex - PPC, 95% CI [2.38, 60.5]; paraventricular nucleus of the thalamus - PVT, 99% CI [9.02, 98.7]; and nucleus reuniens - RE, 99% CI [6.63, 92.8]). Conversely, decreases in hub scores were detected in the intermediate retrosplenial cortex (iRSP, 95% CI [-78.9, 17.6]) and the posterior anterior cingulate cortex (pACC, 95% CI [−79.0, −8.25]).

These findings highlight a shift in the relative influence of specific structures within the network, despite the fact that the overall cohesion of brain connectivity remained unchanged. Reactivation of spatial memory ensembles increased the influence of key nodes, including the dorsal hippocampus, associative cortical regions, and thalamic areas, while reducing the influence of intermediate retrosplenial and posterior anterior cingulate cortices.

### The spatial memory network is governed by a subset of influential structures

To visualize the functional networks of Wt and TetTag mice during peak DCZ (T4), individual networks were averaged. Correlations that were significant (p < 0.001) in at least 75% of the individual networks per group were retained and averaged, ensuring the inclusion of only consistent edges. Communities within these networks were identified using the Louvain algorithm (Blondel et al., 2008), with node hub scores represented by fill color and size, and edge weights represented by color and thickness (Figure 5A).

**Figure 5.**
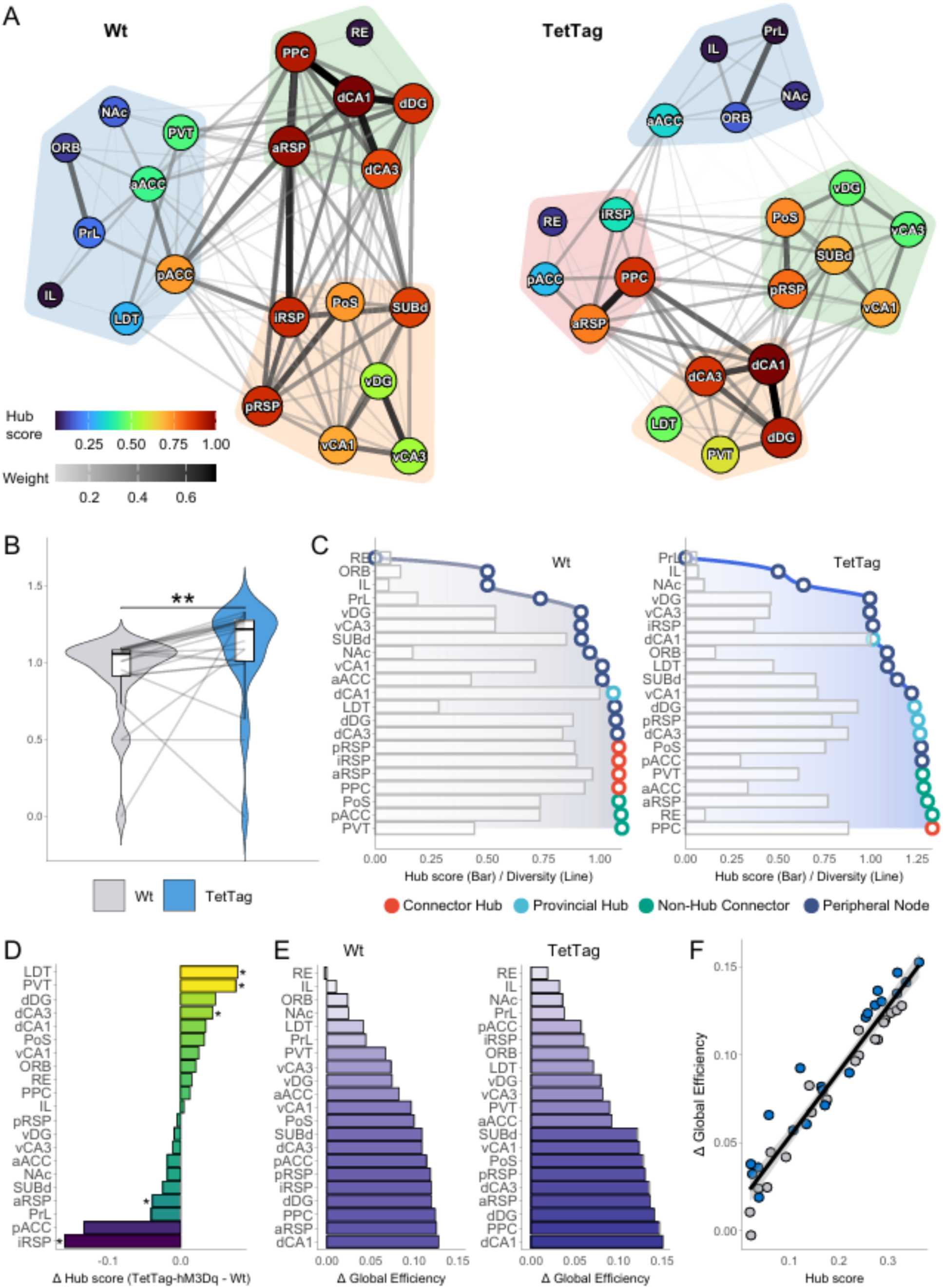
**Spatial memory network is modular and governed by a subset of influential structures** A) Graph visualization of the network topology of Wt (left) and TetTag (right) groups. B) Comparison of connection diversity between groups. Asterisks denote Mann-Whitney’ s test statistical significance, ** p < 0.01. C) Classification of node roles in Wt (left) and TetTag (right) networks based on hub scores (bars) and diversity scores (lines). Points are color-coded to indicate roles: Connector Hubs, Provincial Hubs, Non-Hub Connectors, and Peripheral Nodes. D) Differences in hub scores between TetTag and Wt nodes. Asterisks indicate statistically significant differences determined by permutation tests (95% CI). E) Impact of node deletions on global efficiency (ΔGE) assessed using a disruption propagation model in Wt (left) and TetTag (right) networks. E) Pearson’s correlation between Hub scores and ΔGE. N = 8.

The Wt network was divided into three communities (modularity score = 0.16): a dorsal hippocampus cluster including the nucleus reuniens, posterior parietal cortex, and anterior retrosplenial cortex; a ventral hippocampal cluster with the subiculum, intermediate and posterior retrosplenial cortex; and a prefrontal cortex cluster consisting of the orbitofrontal cortex, infralimbic cortex, and prelimbic cortex, anterior and posterior anterior cingulate cortex, nucleus accumbens, laterodorsal thalamus, and paraventricular nucleus of the thalamus(Figure 5A, left). Given that these animals did not undergo ensemble reactivation and were anesthetized, this may reflect resting-state activity (Paasonen et al., 2018).

In contrast, the TetTag network exhibited greater modularity (modularity score = 0.22) with four communities: a dHPC cluster including the LDT and PVT, densely connected to a vHPC cluster containing the subiculum and pRSP; a PFC cluster sparsely connected to the hippocampal clusters, primarily through ORB and NAc; and an additional cluster comprising the pACC, PCC, RE, and a/iRSP, which redistributed PFC inputs to the hippocampal clusters (Figure 5A, right). These differences reveal that the TetTag network displays more localized and modular connectivity compared to controls.

We quantified the diversity of connections in the networks by calculating the Shannon entropy of the communities reached by each node’s neighbors (Figure 5B). This measure reflects how broadly a node connects across different communities. Although the networks did not differ regarding clustering, efficiency and edge density (Figure 4A), nodes in the TetTag network had significantly higher diversity compared to Wt nodes (Mann-Whitney’s test, p = 0.006). This indicates that memory ensemble reactivation enhances cross-community connectivity.

Next, we assessed the diversity and hub score values of individual structures across networks (Figure 5C). Bars represent hub scores, while points indicate diversity values. Within each group, structures were classified based on these scores: those in the top 20% of hub scores were identified as hubs, and those in the top 20% of diversity values were designated as connectors. Structures falling into both categories were classified as Connector Hubs, while those only among the top hub score structures were labeled as Provincial Hubs. Structures not classified as hubs were categorized as Non-Hub Connectors if they scored high in diversity, or otherwise were categorized as Peripheral Nodes. In both networks, dCA1 emerged as the top hub region but was not a connector, indicating its influence is largely confined to the dHPC community. In the Wt network, the PCC and all RSP subregions were classified as Connector Hubs, while in the TetTag network, only the PCC retained its role as a Connector Hub, with the highest diversity score. The aRSP in TetTag was categorized as a Non-Hub Connector, and other RSP subregions were identified as Peripheral Nodes. Notably, in the TetTag network, non-hub regions such as the PVT, aACC, and RE acted as connectors, while regions like dDG, dCA3, and pRSP served as Provincial Hubs. In contrast, only dCA1 was a Provincial Hub in the Wt network. Interestingly, the RE ranked last in diversity in the Wt network but rose to second place during engram reactivation. These changes indicate a sharper division of roles in the engram-reactivation network, likely reflecting an enhanced ability to balance functional specialization within distinct modules with effective inter-community communication. This higher diversity, combined with specialized hubs and connectors, may underlie the network’s capacity to sustain memory-guided behavior.

To highlight differences in node importance induced by memory ensemble reactivation, we calculated differences in hub score between networks (Figure 5D) using a permutation-based approach with network rewiring (10,000 iterations). It revealed significant increases in hub scores for LD, PVT, and dCA3 compared to controls. Conversely, hub scores decreased in the iRSP and aRSP. We further assessed the importance of individual nodes to network efficiency by simulating node deletions using a disruption propagation model (Figure 5E) (Vetere et al., 2017). In both groups, deletion of dCA1 caused the largest drop in global efficiency. However, deletions of pACC and iRSP had a reduced impact in the TetTag network, consistent with hub score differences. While the Wt network showed a gradual change in node deletion impacts, the TetTag network exhibited a clear separation between high-impact and low-impact nodes, further indicating a more specialized network structure. (Heuvel and Sporns, 2011). A strong correlation between hub scores and efficiency impact was observed (Pearson’s r = 0.96, p < 0.001; Figure 5F), highlighting the predictive power of eigencentrality hub score metric for nodal importance. These results reveal that neurons active during spatial memory retrieval are sufficient to rewire network-wide connectivity patterns related to the functional specialization of individual brain areas.

### Behavior-Predictive Hubs form A Minimal Network centered around the dHPC

Although several brain areas increased their network influence following memory ensemble reactivation, not necessarily all were linked to memory retrieval, or task performance. To identify areas whose increased influence was associated with memory recall, we examined the correlation between hub scores and behavioral performance during the late EL phase (Figure 6A). At that stage of the T-Maze task, control animals exhibited reduced correct responses, whereas TetTag mice maintained the learned preference for the previously rewarded maze arm (Figure 2A).

**Figure 6.**
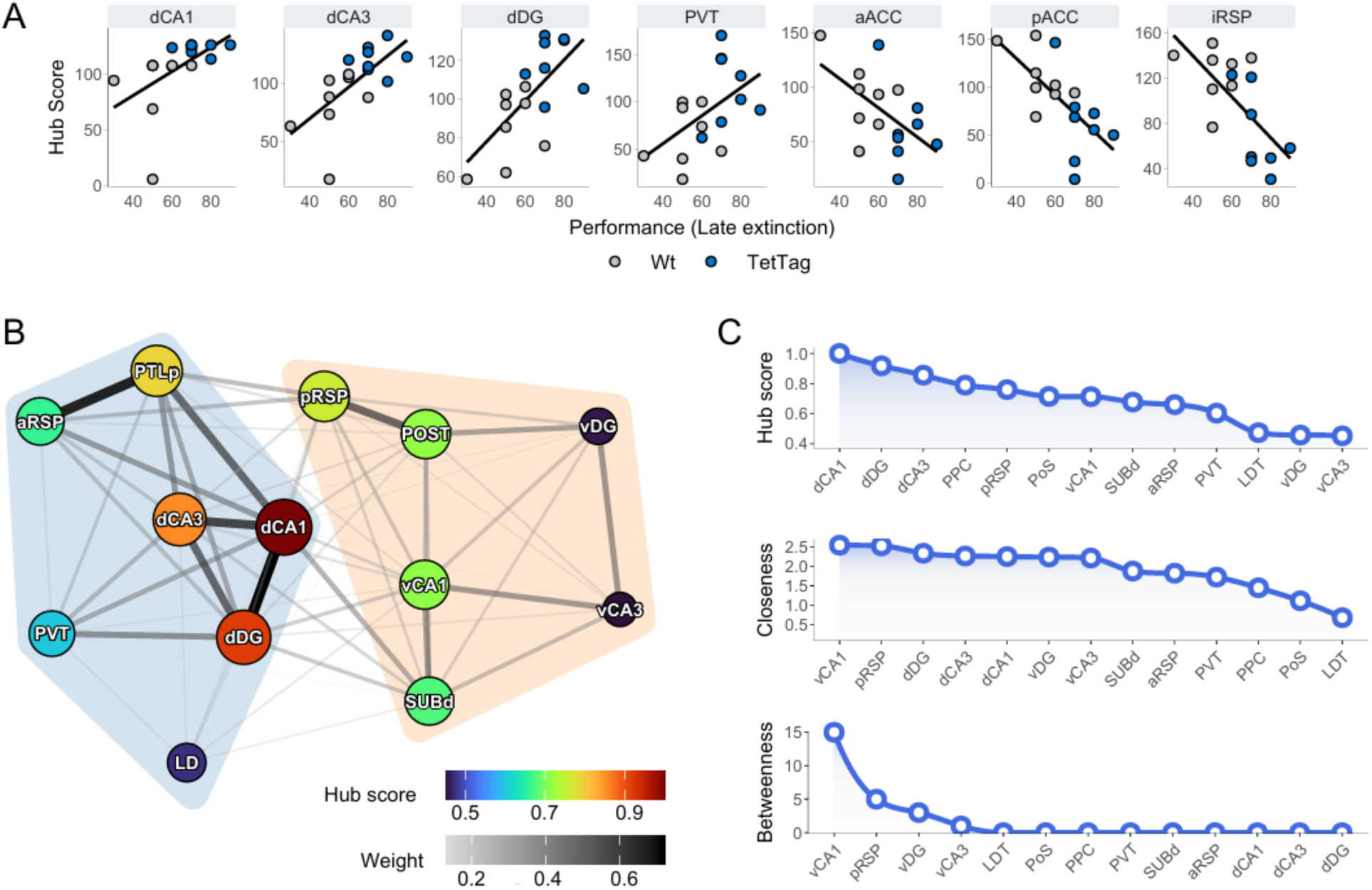
**Behavior-Associated Hubs and Minimal Memory Network.** A) Correlation of hub scores with behavioral performance during the last extinction block, identifying seven regions with significant Pearson’s correlations. B) Minimal memory network constructed from regions with positive, significant correlations between hub scores and performance, along with their immediate connections. Node hub scores and edge weights are color-scaled. C) Rankings of hub scores, closeness centrality, and betweenness centrality values for the minimal memory network.

Hub scores of dCA1 (r = 0.52, p = 0.034), dCA3 (r = 0.66, p = 0.005), dDG (r = 0.68, p < 0.005), and PVT (r = 0.52, p = 0.04) showed positive Pearson’s correlations with behavioral performance, in accordance with the critical role of these regions in spatial memories. Conversely, aACC (r = −0.55, p = 0.03), pACC (r = −0.67, p = 0.005) and iRSP (r = −0.67, p = 0.005) exhibited negative correlations, consistent with their reduced influence in the TetTag network (Figures 4B, 5B).

To explore the subnetwork associated with memory-guided behavior, we constructed a minimal TetTag network comprising the regions with hub scores positively correlated with late extinction performance and their immediate connections, and communities within these networks were identified using the Louvain algorithm (Blondel et al., 2008) (Figure 6B). This network revealed two main communities: a dHPC-centered community (including dCA1, dCA3, dDG, LDT, PVT, PCC, and aRSP) and a vHPC-centered community (consisting of vCA3, SUBd, POST, and pRSP). Within this framework, dCA1, dCA3, and dDG emerged as the most influential hubs, driving network integration, while vDG and vCA3 exhibited the lowest influence (Figure 5C).

Notably, despite their lower hub scores compared to dHPC regions, vCA1 and pRSP ranked highest in closeness and betweenness centrality, indicating a role in facilitating information flow within and across communities (Figure 6C).

While this minimal network does not capture the full complexity of relationships in the broader network (Figure 5), it highlights the local connectivity patterns of nodes most closely linked to memory-related behavior. Consistent with their complementary functions (Contreras et al., 2018; Lee et al., 2019; Ramos and Morón, 2022), these findings reveal that while the dHPC is the minimal network’s core component, it maintains close functional relationships with the ventral hippocampus, relying on critical connectors and modulatory inputs from the subiculum, thalamus, and retrosplenial cortex.

## DISCUSSION

Animals possess the ability to learn rewarded locations and to use spatial cues to efficiently navigate toward them. Equally important is their ability to adaptively modify their strategies during extinction learning (EL) when contingencies change. These memory processes are thought to be mediated by sparse populations of neurons, embedded within interconnected brain circuits (Josselyn and Frankland, 2018; Milczarek et al., 2018). However, knowledge about how these memory ensembles influence interregional connectivity is limited. To address this, we used TetTag-hM3Dq mice to insert the excitatory hM3Dq receptor into neurons that were activated during appetitive spatial learning. We then chemogenetically reactivated the tagged neuronal, first during EL and later during fMRI, to examine which brain networks support a successfully learned spatial appetitive task. We demonstrated that the reactivation of neuronal ensembles that were active during successful task acquisition, significantly disrupts EL, given that animals persisted in visiting the target arm, even though a reward was no longer present. At the network level, ensemble reactivation promoted modularity and nodal specialization, reconfiguring hub regions by increasing the importance of some nodes, while reducing that of others. We identified a network “backbone”, closely associated with spatial memory expression, that extends beyond the dorsal hippocampus. This backbone includes ventral CA1, the subiculum and postsubiculum, the posterior parietal cortex, and the retrosplenial cortices, along with the paraventricular and laterodorsal thalamic relays. These findings indicate that neuronal ensembles that are engaged during spatial learning drive goal-directed behaviors by orchestrating connectivity patterns, which guide brain-wide information flow through a cortico-hippocampal-thalamic circuit, the activation of which impedes EL.

Previous studies, using gain- or loss-of-function approaches, have shown that reactivating or silencing neurons that are active during memory encoding, or retrieval, can respectively promote or inhibit learned responses (Frankland et al., 2019). These findings have been instrumental in identifying circuits within specific brain structures that are sufficient and/or necessary for various types of memories. For example, neuronal activation or silencing in the hippocampus (Denny et al., 2014; Liu et al., 2012) and amygdala (Gore et al., 2015) have been shown to either enhance or diminish fear-conditioned responses in rodents. In this study, we used TetTag-hM3Dq mice to identify neurons across the brain that were active when animals successfully navigated a T-maze to find a low-probability reward. Tagging was performed at a training stage when animals had already reached peak task acquisition performance, therefore capturing both the retrieval and reinstatement of a well-learned spatial memory, rather than tagging neurons that were involved in *de novo* experience encoding. This strategy revealed the participation of sparse populations of neurons across the brain. We then tested the effect of ensemble reactivation driven by DCZ treatment during EL in the unchanged T-maze context. In mice, successful EL in an AAA paradigm, as used here, typically requires two days of exposure to the unrewarded T-maze (Méndez-Couz et al., 2021). This timeline was also observed in the control group in the present study, in which DCZ had no effect on neuronal ensembles due to the absence of neuronal tagging and the pharmacological selectivity for hM3Dq receptors (Nagai et al., 2020). However, when tagged ensembles were reactivated with DCZ, EL was prevented, and animals continued to show behavior consistent with the task acquired on the acquisition days. These results suggest that the tagged ensembles prompted memory reactivation and perseverance in the previously learned task, though the reward was absent. They also suggest that the reactivation of these ensembles is associated with EL failure.

To investigate the role of these ensembles upon brain wide function, the tagged neurons were reactivated by DCZ during fMRI. Memory ensemble reactivation did not induce abrupt changes in the overall distribution of, or mean values of interregional correlation coefficients in, the BOLD signal compared to control animals. This overall stability is not unexpected under anesthesia, which is known to suppress neural activity and interregional connectivity (Grandjean et al., 2014; Wu et al., 2016). Moreover, the anesthesia-induced reduction in background activity enhances signal-to-noise ratios (Grandjean et al., 2014; Peeters et al., 2001), likely highlighting specific effects driven by hM3Dq receptor activation against a suppressed baseline. Indeed, specific changes emerged between pairs of brain regions. To better understand these alterations, we applied graph theory network analysis (Haubrich and Nader, 2023; Sporns, 2018). Metrics such as network efficiency, density, and clustering did not differ significantly following hM3Dq receptor activation, indicating that average properties across the whole network were not affected. However, we observed an increase in average eigenvector centrality values, which reflect the influence of nodes within the network (Lohmann et al., 2010). In other words, specific brain regions became dominant drivers of the networks. Given that higher eigenvector centrality reflects that nodes connect to other nodes that are themselves highly connected (Fagerholm et al., 2015), this indicated the emergence of prominent hub structures that played a role in memory retrieval and reinstatement.

The prominent hub structures identified included the dorsal hippocampus subfields (dDG, dCA3, dCA1), paraventricular (PVT) and reuniens (RE) thalamic nuclei, and posterior parietal association area (PCC). In contrast, the intermediate segment of the retrosplenial (iRSP) and posterior segment of the anterior cingulate cortex (pACC) was less influential. The central influence of dHPC was expected, given its well-established role in memory and navigation (Hagena and Manahan-Vaughan, 2024; Lisman et al., 2017; Stacho and Manahan-Vaughan, 2022a). Similarly, the PCC’s central importance as a connector hub aligns with its involvement in spatial processing (Kravitz et al., 2011). The RSP, critical for dynamically integrating trajectories within a what-when-where framework (Stacho and Manahan-Vaughan, 2022b), exhibited a region-specific shift: iRSP had only peripheral network importance, while the anterior and posterior segments acted as connectors and hubs in the network, suggesting functional specialization within the RSP. The PVT and RE also emerged as key players. The PVT, important for relaying homeostatic information to hippocampal and cortical regions (Penzo and Gao, 2021), and the RE, involved in relaying spatial information between the hippocampus and subcortical regions (Griffin, 2021), both demonstrated network prominence as important connector areas. These connectivity patterns reveal that the activation of neurons engaged in spatial memory retrieval and reinstatement reshapes brain network connectivity, causing a subset of regions to become important hubs. This finding compelled us to further scrutinize how these changes affect network organization and to delineate the specific roles of each region within the network.

The balance between functional segregation and integration across a network is important for maintaining specialized processing within distinct groups of brain regions while also allowing for the exchange of information between them (Wig, 2017). For instance, in humans, balanced network integration and segregation is particularly relevant for memory above other cognitive functions (Wang et al., 2021). To explore these dynamics, we built network models and looked for groups of tightly interconnected regions, known as communities, that work together to process information, enabling segregated information processing (Deco et al., 2015; Sporns, 2013). Reactivating memory ensembles increased network modularity, with brain regions clustering into smaller communities, while also broadening cross-community connectivity. This aligns with human fMRI studies demonstrating that, in contrast to a motor task, cross-community connectivity is crucial for memory task performance (Cohen and D’Esposito, 2016; Keerativittayayut et al., 2018). Based on their intra- and cross-community links, we also classified brain areas according to their functional roles in the networks (Guimerà and Nunes Amaral, 2005) and found that the TetTag network exhibited a sharper division of roles: more regions were classified as local hubs or non-hub connectors, and fewer as both connectors and hubs, compared to controls. Together, these findings reveal that memory ensemble reactivation is sufficient to promote functional specialization, while maintaining effective inter-community communication through relay regions.

Notably, a network community comprising the pACC, RE, a/iRSP, and PCC emerged as a critical intermediary for information flow between the prefrontal cortex and dHPC. Given the established roles of the ACC (Frankland and Bontempi, 2005), RE (Aggleton and O’Mara, 2022), RSP (Stacho and Manahan-Vaughan, 2022b), and PCC (Kravitz et al., 2011) in multimodal sensory integration and memory, the emergence of this community may reflect the integration of mnemonic and spatial signals from the dHPC with executive processing from the PFC. Additionally, the LDT and PVT moved closer to the dHPC fields compared to controls, in line with their contribution to increased relay of sensory information into the hippocampus during memory acquisition (Roy et al., 2022). On the other hand, the ORB and NAc became the primary gateway linking the PFC and vHPC. The OFC is known for decision-making and updating associations (Lissek et al., 2020; Padoa-Schioppa and Conen, 2017; Ranganath and Ritchey, 2012), while the NAc links environmental cues with rewarding outcomes (Ito et al., 2008; Trouche et al., 2019). This mediation may reflect a mechanism by which motivational signals integrate with executive control to drive goal-oriented processing in the vHPC.

In essence, these findings indicate that reactivating spatial memory neurons reorganizes the brain network around a set of specialized hubs distributed across distinct communities and interconnected by relay regions. To identify the backbone of this core configuration, we applied a disruption propagation model (Vetere et al., 2017) to systematically delete nodes and thereby assess their impact on network efficiency (Tu et al., 2021). While nodal disruptions strongly correlated with hub scores in both groups, in the control network the effects of node deletion were distributed along a continuum, reflecting a state of low specialization under anesthesia. In contrast, the TetTag network demonstrated a concentrated reliance on a specific group of regions. As expected, the dorsal hippocampal subfields were part of the network backbone. Beyond the well-established importance of the dHPC, we also found that the other key members of the network were the vCA1, the subiculum and postsubiculum, the posterior parietal cortex (PCC), and the anterior and posterior segments of the retrosplenial cortex. This pattern suggests that information flow across the network is heavily dependent on a dHPC–vCA1–SUB–PoS– PCC–RSP backbone.

The wider network included structures which may reflect the engagement of neurons linked to both pure spatial memory recall and updating (dHPC), as well as other functions crucial for goal-directed navigation, such as decision-making (PFC) and the processing of sensory cues, and motivation (O’Doherty et al., 2017; Sosa and Giocomo, 2021). To identify regions where network influence was tied to task performance, we correlated hub scores with late EL performance - a period during which only TetTag mice exhibited sustained behavior. Positive correlations were found for dHPC subfields and PVT, while negative correlations were observed for ACC and iRSP. These results suggest that increased network influence by the dHPC circuit is associated with the behavioral expression of goal-directed, appetitive spatial memory with thalamic participation. To reveal the connectivity patterns driving this outcome, we generated a minimal network containing the dHPC regions and their direct connections. The resulting minimal network contained all regions previously identified as forming the TetTag’s network backbone, and additional thalamic structures. Clustering analysis revealed two distinct communities: a dHPC-centered cluster and a vHPC-centered cluster. Within the dHPC cluster, the dCA1-dCA3-dDG subcircuit exhibited the highest network influence, with neighboring nodes including PVT, LDT, aRSP, and PCC. The vHPC cluster contained SUBd, pRSP, and PoS, with vCA3 and vDG showing the least influence. Although the dHPC subfields dominated in terms of hub values, the vCA1 and pRSP ranked highest in closeness and betweenness and were therefore central nodes for transmitting information within and between communities. This suggests that while dHPC regions serve as primary hubs, connector nodes, such as vCA1 and pRSP, play crucial roles in maintaining the network’s functional coherence.

Our finding that the ventral hippocampus was involved in spatial memory acquisition is consistent with reports by others (Beer et al., 2014; Lee et al., 2019; Ramos, 2022). Interactions of the dorsal and ventral hippocampus during spatial navigation have been demonstrated (Lee et al., 2019), whereby the ventral hippocampus may be especially important for post-learning memory consolidation (Ramos, 2022). The ventral hippocampus sends afferents to the prefrontal cortex (PFC) (Liu and Carter, 2018) and disruption of this communication disrupts cognitive flexibility and spatial working memory (Blot et al., 2015). Successful spatial learning requires bidirectional information flow between ventral hippocampus and PFC (Xia et al., 2019) and spatial information updating requires interaction between the two structures (Park et al., 2021). Interestingly we did not detect direct connectivity between the ventral hippocampus and the infralimbic and prelimbic cortices during ensemble reactivation, although afferents to these regions have been described (Liu and Carter, 2018). This may be because these inputs are more relevant to EL (Brockway et al., 2023) and fear memory (Twining et al., 2020). In addition, when contrasting how influential brain areas were in the TetTag network versus the control network, the thalamic LDT and PVT, along with dCA3, exhibited the most significant positive shifts in network influence upon memory ensembles activation. This likely supported persistent goal-directed behavior during EL, as connectivity among these regions is critical for mnemonic biasing of attention (de Bourbon-Teles et al., 2014).

In conclusion, the results of this study show that sparse yet broadly distributed neuronal ensembles that are activated during appetitive spatial memory retrieval and reinstatement are sufficient to reinforce learned responses and inhibit EL. These ensembles drive significant network reorganization, enhancing the brain’s modular specialization while preserving cross-community integration through emergent hubs and gateways. Although the dorsal hippocampus remains central to information flow, effective network communication also critically depends on ventral CA1, the retrosplenial cortex, the posterior parietal cortex, and thalamic nuclei - together forming a core circuit that supports goal-directed navigation and counteracts EL. Furthermore, top-down prefrontal control over the dorsal hippocampus is mediated via a PPC-RSP-RE-ACC circuit, whereas control over the ventral hippocampus implicates the NAc and ORB. In effect, the interference of EL was caused by the emulation of reactivation of well-established recent memory, indicating that memory decay (of the previously learned experience) may be an important facet of effective EL In sum. we propose that spatial memory-associated ensembles (Denny et al., 2017) support memory retrieval and reinstatement by inducing widespread changes in interregional communication, channeling information through key hub and modulatory regions that extend beyond the dorsal hippocampus. The dominance of these hubs is sufficient to override EL despite changes in behavioral contingencies.

## ACKNOWLEDGMENTS

We are grateful to Jens Colitti-Klausnitzer and Juliane Böge for technical assistance with animal treatments, Nadine Kollosch for animal care, Beate Krenzek for genotyping, Dr. Thu-Huong Hoang and student assistant, Rabianur Kartalcik, for support with immunohistochemistry, and Xavier Helluy for assistance with fMRI. This work was supported by a German Research Foundation (Deutsche Forschungsgemeinschaft, DFG) grant to DM-V (SFB 1280/A04, project number: 316803389). The authors declare no conflict of interest.

