## Supplementary Figures 1 and 2 for "Sparse memory ensembles set brain-wide network states to sustain learned associations"

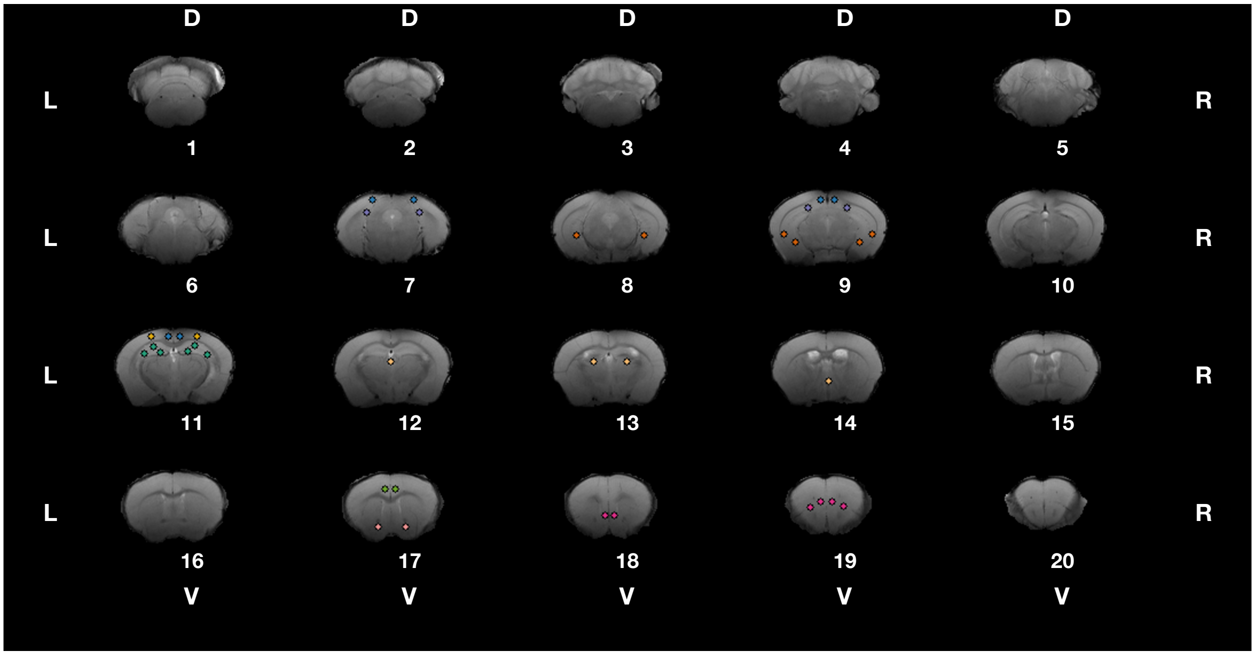


**Supplementary Figure 1**

Coronal slices processed during fMRI. L, R, D, and V denote the left, right, dorsal, and ventral directions, respectively. Slices are numbered from posterior to anterior. Seeds are marked with colored circles.

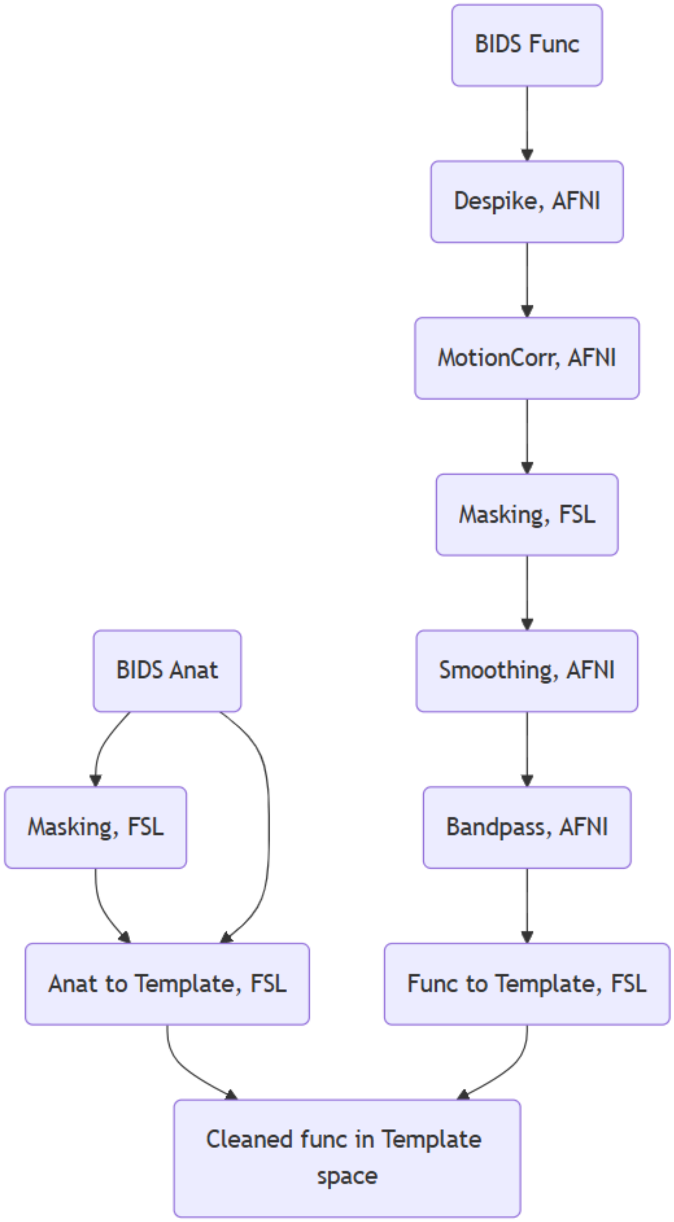


**Supplementary Figure 2**

Diagram of the fMRI pipeline for data pre-processing and time-series extraction.
